# Microplastic fibers affect dynamics and intensity of CO_2_ and N_2_O fluxes from soil differently

**DOI:** 10.1101/2020.09.21.306589

**Authors:** Matthias C. Rillig, Mathias Hoffmann, Anika Lehmann, Yun Liang, Matthias Lück, Jürgen Augustin

**Affiliations:** Freie Universität Berlin, Institut für Biologie, Altensteinstr. 6, D-14195 Berlin, Germany; Berlin-Brandenburg Institute of Advanced Biodiversity Research (BBIB), D-14195 Berlin, Germany; Leibniz Centre for Agricultural Landscape Research (ZALF), Research Area 1 “Landscape Functioning”, Eberswalder Str. 84, D-15374 Müncheberg, Germany

**Keywords:** Nitrous oxide, carbon dioxide, soil structure, microplastic fibers, greenhouse gas

## Abstract

Microplastics may affect soil ecosystem functioning in critical ways, with previously documented effects including changes in soil structure and water dynamics; this suggests that microbial populations and the processes they mediate could also be affected. Given the importance for global carbon and nitrogen cycle and greenhouse warming potential, we here experimentally examined potential effects of plastic microfiber additions on CO_2_ and N_2_O greenhouse gas fluxes. We carried out a fully factorial laboratory experiment with the factors presence of microplastic fibers (0.4% w/w) and addition of urea fertilizer (100 mg N kg^−1^). The conditions in an intensively N-fertilized arable soil were simulated by adding biogas digestate at the beginning of the incubation to all samples. We continuously monitored CO_2_ and N_2_O emissions from soil before and after urea application using a custom-built flow-through steady-state system, and we assessed soil properties, including soil structure. Microplastics affected soil properties, notably increasing soil aggregate water-stability and pneumatic conductivity, and caused changes in the dynamics and overall level of emission of both gases, but in opposite directions: overall fluxes of CO_2_ were increased by microplastic presence, whereas N_2_O emission were decreased, a pattern that was intensified following urea addition. This divergent response is explained by effects of microplastic on soil structure, with the increased air permeability likely improving O_2_ supply: this will have stimulated CO_2_ production, since mineralization benefits from better aeration. Increased O_2_ would at the same time have inhibited denitrification, a process contributing to N_2_O emissions, thus likely explaining the decrease in the latter. Our results clearly suggest that microplastic consequences for greenhouse gas emissions should become an integral part of future impact assessments, and that to understand such responses, soil structure should be assessed.

## Introduction

As a result of human activities, the load of reactive nitrogen compounds (NH_3_/NH_4_, NO_3_^−^, NO_x_, N_2_O) on the earth has more than doubled in recent decades ^1, 2^. This was accompanied by a doubling of the intensity of the global nitrogen cycle. A main driver of this development are intensified agricultural practices entailing increased application of synthetic nitrogen fertilizers since the end of the Second World War ^3-5^. While ensuring food security for an ever-growing world population ^6^, agriculture has developed globally into a major source of climate-relevant trace gases. This applies in particular to nitrous oxide. Agriculture accounts for 60% of the total man-made nitrous oxide release ^7^. The continuing increase in N fertilization or N surplus in production also appears to be the main reason for the unexpectedly strong acceleration in atmospheric N_2_0 concentration in recent times ^8, 9^. Consequences for the role of soils as sources and sinks of the important greenhouse gas CO_2_ can also be expected, since the carbon and nitrogen budgets of soils are closely linked. This is especially true for the mineralization of soil organic matter as a source of CO_2_ release from soils ^10^. A prerequisite for the development of effective strategies to reduce land-use-related greenhouse gas emissions is comprehensive knowledge of the relevant processes and their regulation by internal and external drivers ^11-13^. An important process that has been under-researched in microplastic-affected soils is the emission of greenhouse gases ^14^. Despite its potential importance, compared to other factors of global change, we have so far only scratched the surface in terms of assessing microplastic impacts on soil properties and processes in general ^15-18^.

Microplastic pollution is becoming increasingly recognized as a factor of global change, affecting not only aquatic but also terrestrial ecosystems and the soil ^17, 19^. Microplastics occur as primary microplastic or secondary microplastic and in a wide variety of sizes, shapes, chemistries and with a huge diversity of additives. Microplastic particles are expected to arrive in most ecosystems via aerial deposition ^20, 21^, but in agroecosystems there are also other input pathways including addition of sewage sludge or compost, which have been estimated to represent rather large input fluxes ^16^. Once they have arrived in agroecosystems, there are a range of plausible pathways (including plowing) that lead to a transport of such particles into the soil ^22^, where effects upon soil properties, processes and biodiversity can then unfold. Previous studies on the effects of microplastics have shown effects on soil organisms, especially microorganisms, and chemical conversion processes in soils ^23-25^. Initial evidence also pointed to soil physical properties being altered by microplastics ^26^. We have evidence that microplastic can affect basic parameters including soil structure and bulk density ^15^, and that the performance of biota can be altered, which has been shown, for example for earthworms ^27^, microbes ^28^, and for plant growth ^29-32^. However, the only study to date on the effect of microplastics on the emission of climate-relevant trace gases from soils does not address the impact of soil physical properties on the greenhouse gas emission ^28^.

However, the fact that the intensity of N_2_O and CO_2_ release is very strongly determined by soil physical properties, irrespective of the amount of N fertilization, has been shown in numerous studies. Parameters of the soil structure such as air permeability, aggregate size distribution, and size and design of pore space seem to play an important role. On the one hand, they have a direct influence on the movement of the gases via mass flow and diffusion in the soil in a variety of ways and thus also the emission of greenhouse gases ^33-41^. On the other hand, they act indirectly by controlling the availability of oxygen in the soil, which in turn influences the processes of CO_2_ and N_2_O formation and N_2_O consumption in many ways ^42-45^.

In view of this, it is quite possible that the changes in soil structure caused by microplastics could indeed have an impact on the release of climate-relevant trace gases. The aim of our investigations here was thus to contribute to the clarification of the effect of microplastic addition on the emission of the greenhouse gases CO_2_ and N_2_O in interaction with N fertilization. We wished to test two main hypotheses: (i) addition of microplastics leads to significant changes in important soil physical properties, most notably soil aggregation; and (ii) effect of N fertilization on the release of N20 and CO2 are therefore altered by the addition of microplastics.

## Materials and Methods

### Soil material

The soil material investigated was taken from the Ap horizon of a non-eroded Albic Luvisol (Cutanic, soil classification according to IUSS Working Group WRB, 2015), consisting of 59% sand, 32% silt, and 9% clay. This site is located on the flat summit within the CarboZALF experimental field, where studies on the influence of erosion on the C-dynamics were conducted. The CARBOZALF field belongs to the hummocky landscape within the Uckermark region (northeastern Germany, 53°23’N, 13°47’O, ∼50-60 m a.s.l.) ^46^. Winter wheat was grown there in the year of sampling. The soil material was dried and stored in this form for several weeks prior to the start of the experiment.

### Sample preparation and treatments

The investigations were carried out using soil samples filled into steel cores with a volume of 250 cm^3^ and a bulk density of 1.4 g dry soil cm^−3^. In preparation for incubations, the dry soil was sieved to 2 mm and then moistened to a water-filled pore volume of 48 %. To stimulate the activity of soil microorganisms similar to the conditions in an intensively N fertilized soil, diluted biogas digestate (dry matter 2.5%, pH 8.3) was used for soil moistening. In this way, the substrate was provided with 290 mg total N respectively 170 mg ammonium N per kg dry soil for all treatments at day zero of incubation (Table 1).

**Table 1.**
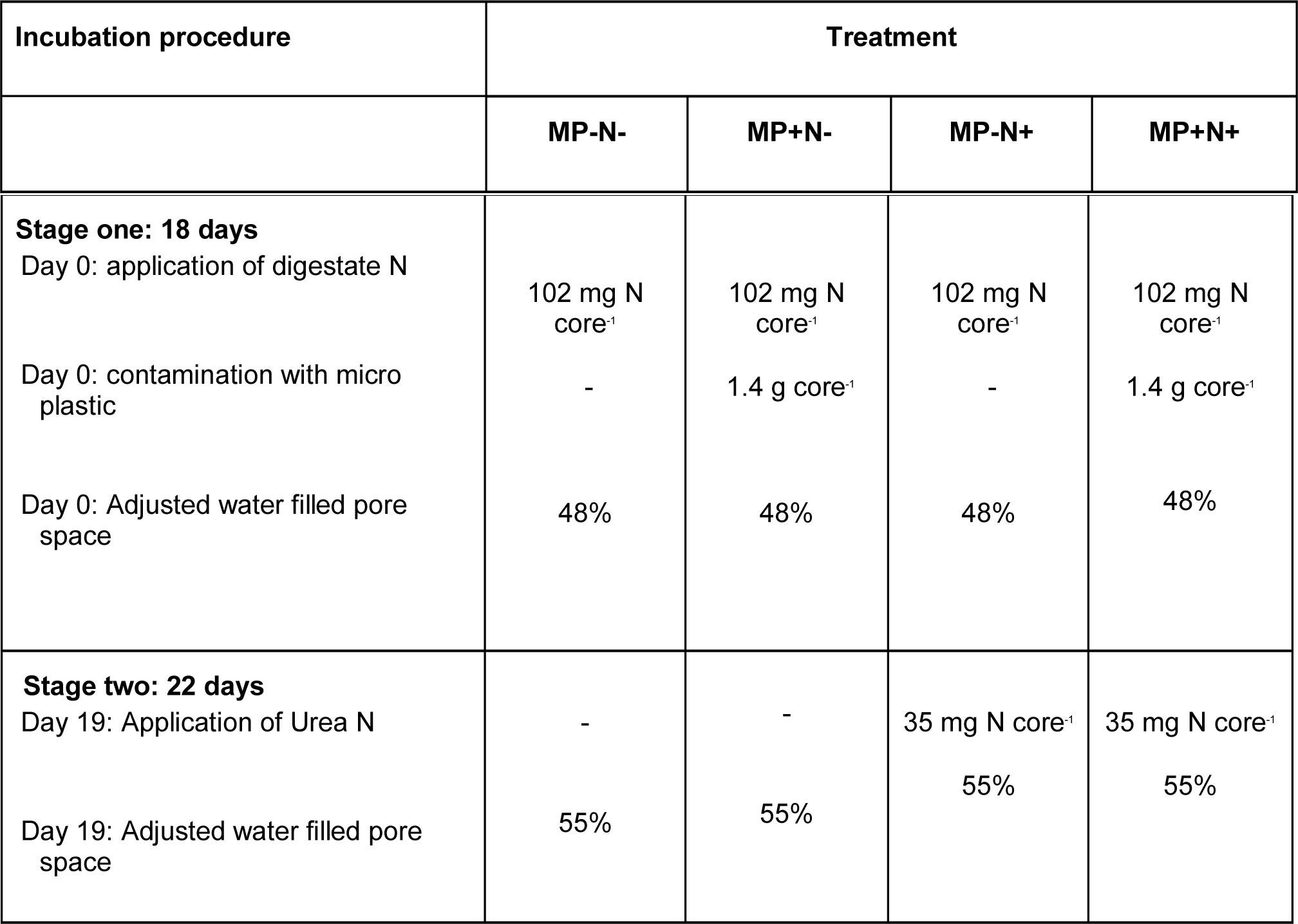
Setup of the two-factorial incubation experiment. Treatment codes are no-addition control (MP-N-), microplastic only (MP+N-), urea only (MP-N+), and addition of both microplastic

To investigate the effect of N fertilization and soil contamination with microplastics alone and in their interaction, four variants were established, each comprising four of these soil cylinders (Table 1). To test the N effect, 35 mg urea-N per core (100 mg N per kg dry soil) was added at day 22 of the incubation.

For contamination of the soil with microplastics we used microplastic fibers, since fibers have repeatedly been shown to affect soil structure ^26, 47-49^, possibly due to their linear shape ^50^. We used polyester fibers (Paraloc rope, 8mm diameter, Mamutec, Switzerland, product number: 0025-00080-01-0), cut by hand (for the size distributions of fiber length and diameter see SI Fig. S1). Fibers were briefly microwaved to minimize microbial loads, following a previous protocol ^30^. The amount of microplastic fibers mixed in was 1.4 g per core (0.4% w/w) at day zero of the incubation.

The microplastic fibers were distributed homogeneously on the surface of the soil substrate. The soil substrate, the diluted biogas digestate and, depending on the variant, also the microfibers were then carefully mixed together and filled into the stainless steel cylinders in layers at day zero of incubation. In each case one treatment without urea fertilization and without microplastic contamination served as control (Table 1). All samples received the same amount of mixing disturbance.

### Incubation experiments

First, the gas emission from the soil cores was monitored over 18 days. On day 19 of incubation, 35 mg urea-N was applied to four samples with or without microplastic contamination. The amount of urea N was dissolved in 10 ml water and injected into the soil using a syringe. Four samples each with and without microplastic contamination were used as controls, into each of which 10 ml of water was injected using a syringe. As a result of this measure the water-filled pore space in all samples increased from 48 to 55%. Subsequently, the gas emissions from the cores was investigated for another 22 days (Table 1).

To determine the CO_2_ and N_2_O emissions, the soil samples were transferred to an incubation facility developed by us (Fig. 1). It works as a flow-through steady-state system corresponding to Livingston and Hutchinson ^51^. The system contains 16 airtight, cylindrical incubation vessels (diameter and height of 13 cm, made from commercially available KG DN sewer pipes and accessories, Marley, Germany), each filled with one soil core. A temperature of 20 °C degrees was maintained in the incubation vessels by means of a climate box. Ambient air flows (32 mL min^−1^) continuously through the headspace of the incubation vessels via channels connecting the pressure vessel and the gas analyzer. In parallel, there is a control channel through which ambient air passes the incubation vessels with the same flow rate directly from the pressure vessel to the gas analyzer. To prevent the soil cores from drying out, the air was saturated to 100% relative humidity before passing through the incubation vessels. Each channel is directly connected to the gas analyzer via a multiplexer and a special circular channel for 7 minutes each. This results in a frequency per measurement and channel of 119 minutes. Gas concentration measurements were performed using cavity ring-down spectroscopy technology in a Picarro G2508 gas concentration analyzer (PICARRO, INC., Santa Clara, USA). Air was circulated between the incubation unit headspace and the CRDS analyzer at 250 mL min^−1^ using a low-leak diaphragm pump (A0702, Picarro, Santa Clara, CA, USA). The air from each of the 17 measuring channels was fed into this circuit via a connecting channel from the multiplexer at a slight overpressure. The overpressure was reduced via another opening on the other side of the circuit, from which the air then flowed out into the environment. In this way the continuous flushing of the circuit with the air from the respective channel to be measured was ensured.

**Figure 1.**
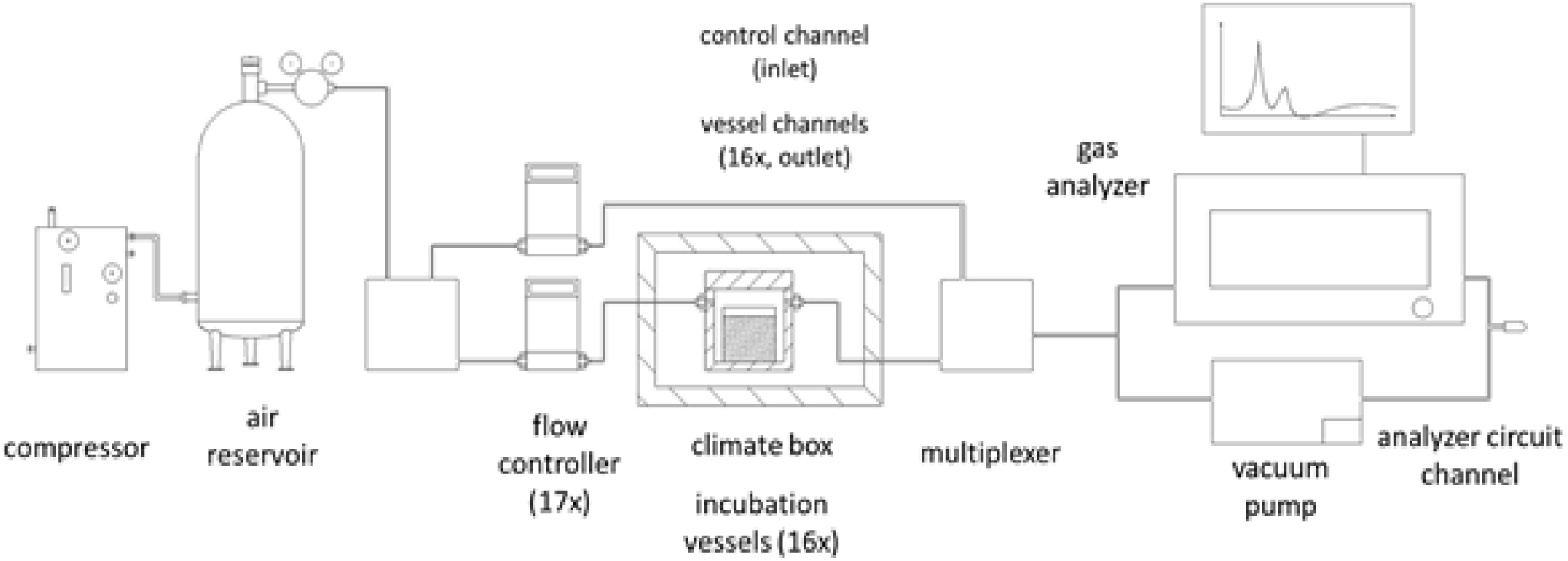
Schematic diagram of the incubation system used to measure CO_2_ and N_2_0 flux rates from the soil cores.

The gas flux rates are calculated from the current gas concentration in the channel, which is connected to the outlet of a specific vessel, and the temporally corresponding concentration in the control channel, which represents the vessel inlet so to say, over time according to Equation 1:

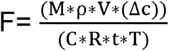

where F is the flux rate (µg CO_2_-C or N_2_O-N core^−1^ h^−1^), M is the molar mass of CO_2_ or N_2_O, respectively (µg mol^−1^), ρ the atmospheric pressure (Pa), V is the air flow rate into the headspace and the channels (m^3^ h^−1^), Δc is the difference of gas concentrations [mol] between outlet of a specific vessel and the control channel, C is the core, R the gas constant (m^3^ Pa K^−1^ mol^−1^), t is the time over which the concentration change was observed, and T the incubation temperature (K).

To avoid bias caused by measurements on the previous channel, only values determined in the last minute of the measurement period of 7 minutes were used to calculate the flux rates. An adaptation of a modular R program script, described in detail by Hoffmann et al. ^52^, was used for the calculation of current gas flux rates and CO_2_-C and N_2_O-N gas losses accumulated over time intervals.

### Soil analysis after incubation

Immediately after the incubation, the air permeability (AP) of the soil cores was measured using the PL-300 device ^53^ based on the Gätke method ^54^. After that, soil subsamples were extracted with 0.0125 M CaCl_2_ solution (ratio 1:4) and analyzed for NH_4_^+^-N and NO_3_^−^-N concentrations using spectrophotometry according to ^55^ with a continuous flow analyzer (Skalar Analytics, CFA-SAN, Breda, Netherlands). Soil moisture content was determined gravimetrically after drying a soil subsample at 105°C until constant weight. Bulk density and water-filled pore space (WFPS) were calculated based on sample volume, dry weight and gravimetric water content. Soil pH was determined in 0.01M CaCl_2_ solution (ratio 1:5) according to DIN ISO 10390. Subsamples were also analyzed for total soil carbon (TC), soil organic carbon (TOC) and total nitrogen (TN) according to DIN ISO 10694 and DIN ISO 13878 using an elemental analyzer (Leco Instruments, TruSpec CNS, St. Joseph, USA). The cold water soluble carbon and nitrogen in the soil was determined by extraction with cold water at the ZALF central laboratory using a Shimadzu TOC-VCPH Carbon Analyzer (Shimadzu Deutschland GmbH, Berlin, Germany).

The measurement of the size class distribution of soil aggregates followed a modified protocol by Kemper and Rosenau ^56^. First, samples were passed carefully through a 4mm sieve. Second, samples were passed through a stack of five sieves (2, 1, 0.25, 0.1 and 0.05 mm) to determine the mass of soil aggregates separating across the resulting six fractions of decreasing particle size. For this, the sieve stack was moved vertically. The movement was kept to a minimum to avoid abrasion but ensure particle separation over the different mesh sizes. The measured mass for each fraction was integrated into the following formula to calculate the mean weight diameter (in mm): 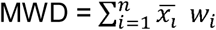 where x_i_ is the mean diameter of size fraction i and w_i_ is the proportion of total soil mass in size fraction i; i.e., soil aggregate size classes are weighted by their mean diameter so that samples with overall larger soil aggregates result in higher MWD values.

Samples were carefully reconstituted and mixed after measuring the MWD before taking 4.0 g of soil. These were placed on a small sieve with 250 µm mesh size, allowed to capilarrily re-wet with deionized water and placed into a sieving machine (Agrisearch Equipment, Eijkelkamp, Giesbeek, Netherlands). During the procedure, the samples were moved vertically for 3 min in metal bins filled with deionized water to experience a disintegrating force. The resulting slaking of the treated soil aggregates caused a separation into a water-stable and water-unstable fraction with a size > 250 µm. From the water-stable fraction, debris (sand particles and organic matter) were extracted to allow calculation of the water-stable aggregate fraction:

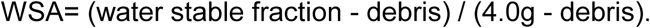

For each sample two technical replicates were tested which were later merged into one mean value for the statistical analysis.

### Statistics

For the statistical analysis, we used a generalized least square model of the “nlme” package ^57^ with implemented varIdent function to account for heterogeneity in the applied treatment (i.e. control, microplastic, urea, microplastic : urea dual application). Control samples were set as reference level. We tested model residuals for normality and heteroscedasticity.

## Results

### Fluxes of CO_2_ and N_2_O

At the beginning of the first phase of incubation an increase in flux rates can be seen for both CO_2_ and N_2_O, followed by a slow decrease. Between the treatments only small differences could be seen (Figure 2). However, at the time of the highest emissions, the variants with microplastics showed slightly lower fluxes for N_2_O and slightly higher fluxes for CO_2_ than the control variants without microplastics. This was accompanied by a differentiated effect of microplastics on the flux dynamics. The presence of microplastics resulted in a faster increase and decrease of flux rates in the case of N_2_O. For CO_2_, the opposite effect was observed. The extreme peak in CO_2_ release occurring in all variants during the first incubation phase is due to system maintenance. The addition of urea at the beginning of the second incubation phase suddenly caused a strong short-term stimulation of the CO_2_ release and a longer-term stimulation of the N_2_O release, whereby the flux rates, especially for N_2_O, were significantly higher than in the first incubation phase (Figure 1). Both gases reacted to the presence of microplastics with the same reaction pattern as in the first incubation phase with regard to the level and dynamics of the emissions. However, the differences between the no-MP addition treatments (MP-N- and MP-N+) and the variants with microplastics (MP+N- and MP+N+) are more pronounced than in the first incubation phase, especially after the emission peak has subsided. In the variants that received only water (MP-N- and MP+N-), the addition of microplastics caused similar behavior in gas emissions. However, the intensity of the reactions was significantly lower than with urea fertilization. Only in the control without microplastics did N_2_O emission after the addition of water reach a similar level as in the first phase of incubation (Figure 1).

**Figure 2.**
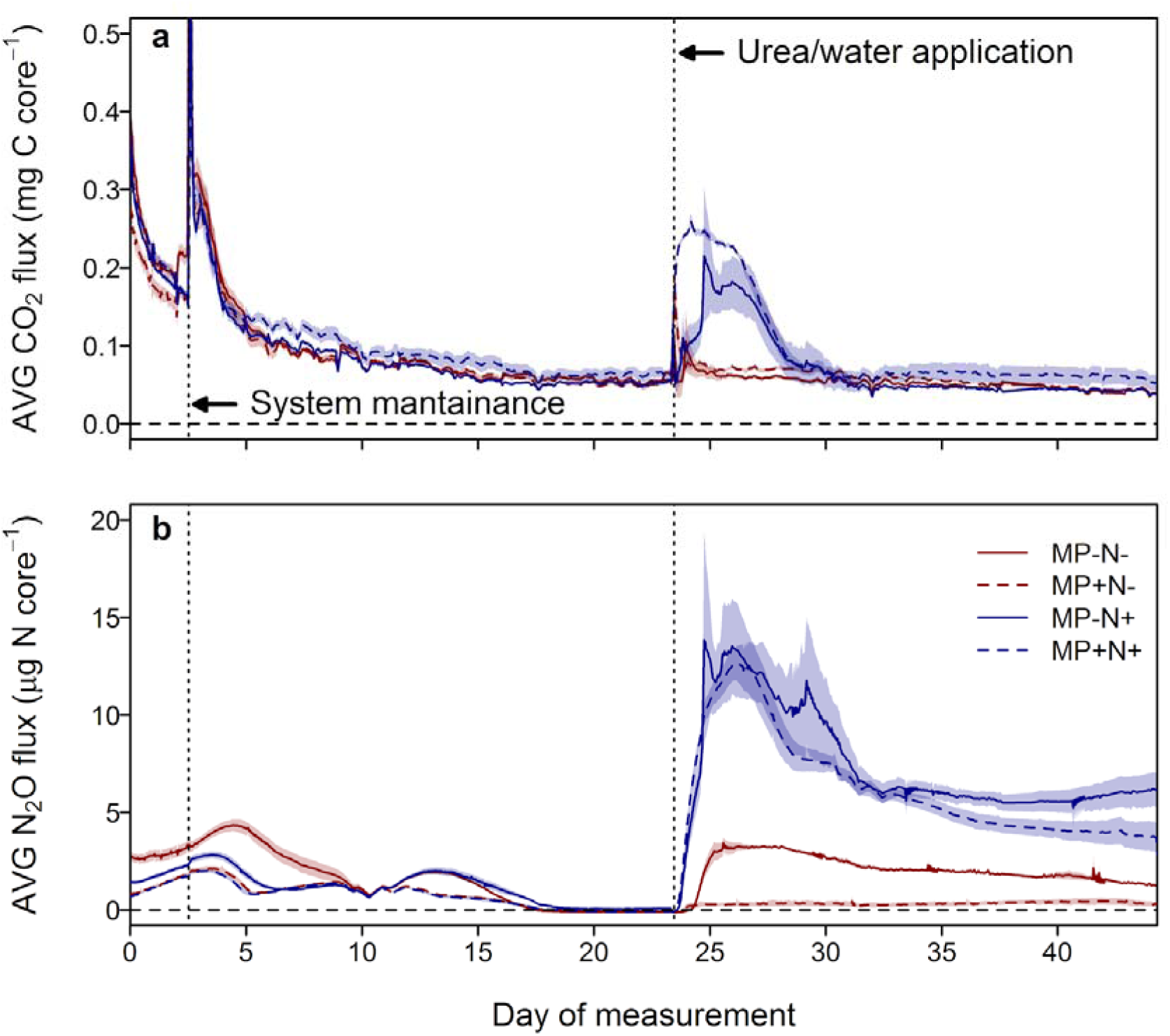
Average (n=4; a) CO_2_ and (b) N_2_O flux dynamics for the four treatments over the time course of the incubation experiment.Treatments are no-addition control (MP-N-; red, solid line), microplastic only (MP+N-; red, dashed line), urea only (MP-N+; blue, solid line), and addition of both microplastic and urea (MP+N+; blue, dashed line). Shaded areas represent ±1 SE.

The effects of microplastics and urea fertilization observed at current flux rates are also reflected in the cumulative CO_2_ and N_2_O emissions (Table 2). Related to the total incubation period, the emitted CO_2_ was slightly promoted by microplastics and strongly by urea application. The strongest effect was caused by the combination of microplastics and urea application. These effects become more apparent when only the second phase of incubation is considered. Looking at the whole period of incubation for N_2_O emission, a clear inhibition of the cumulative release in the presence of microplastics and a weak promotion by urea fertilization becomes apparent. In the second phase of incubation, both the reducing effect of microplastics and the promoting effect of urea fertilization become more evident. Consequently, the same reaction pattern can be seen with regard to the proportion of fertilizer N lost as a result of N_2_O. This proportion was always increased by urea application and reduced by microplastics (Table 1).

**Table 2.**
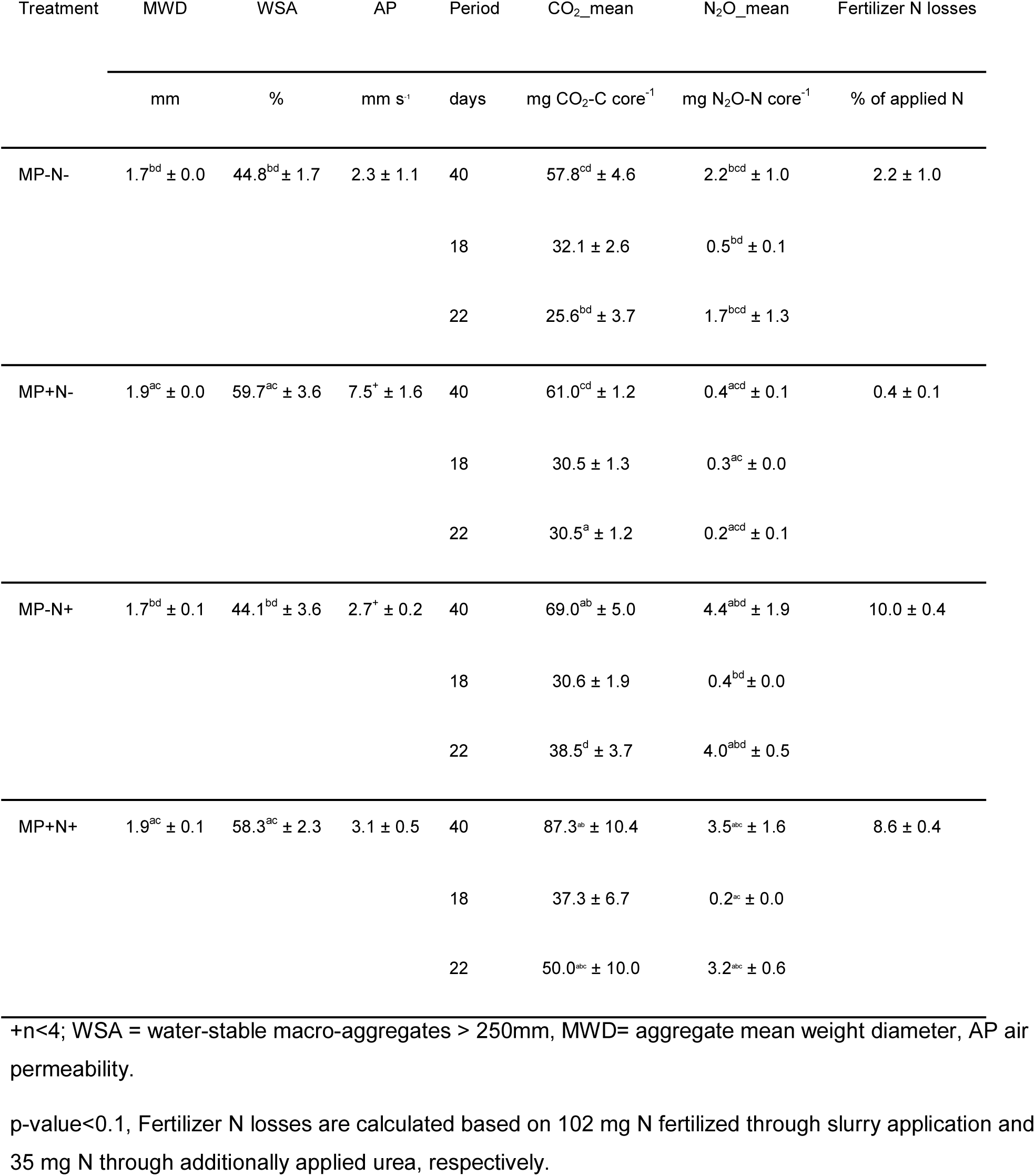
Soil physical characteristics, cumulative CO_2_- and N_2_O emissions and fertilizer N losses by N_2_O

### Physical and chemical soil properties

The two test factors microplastics and urea addition also influenced the physical and chemical soil properties in different ways. The presence of microplastics caused a significant increase in the proportion of water-stable soil aggregates (WSA) and in the mean weight diameter (MWD), and a slight increase in air permeability (Table 2). In contrast, the addition of urea caused a significant increase in the content of cold water soluble carbon and ammonium N in the soil (Table 3, 4). The content of cold water soluble N and of nitrate N was promoted by urea as well as by microplastics. Therefore, the highest values for both parameters again occurred when combining microplastics with urea. The addition of microplastics also resulted in a slight increase in the total and organic carbon content of the soil, likely because microplastic-carbon was co-detected (Table 3, 4).

**Table 3.** Soil chemical characteristics as a function of treatments.

| Treatment |  | pH | TC | TOC | C<sub>CWC</sub> | TN | NH<sub>4</sub>-N | NO<sub>3</sub>-N | N<sub>CWN</sub> |
| --- | --- | --- | --- | --- | --- | --- | --- | --- | --- |
|  |  |  | % soil DM | % soil DM | mg/kg soil DM | % soil DM | mg N kg<sup>-1</sup> soil DM | mg N kg<sup>-1</sup> soil DM | mg kg<sup>-1</sup> soil DM |
| MP-N- | mean | 5.19 | 0.63 | 0.61 | 88.2 | 0.09 | 1.0 | 132.4 | 224.1 |
|  | SD | 0.13 | 0.01 | 0.01 | 4.4 | 0.01 | 0.0 | 6.3 | 2.7 |
| MP+N- | mean | 5.03 | 0.82 | 0.80 | 83.9 | 0.10 | 0.9 | 147.2 | 245.6 |
|  | SD | 0.04 | 0.01 | 0.01 | 13.7 | 0.01 | 0.1 | 1.6 | 5.8 |
| MP-N+ | mean | 5.31 | 0.63 | 0.62 | 142.3 | 0.10 | 20.0 | 144.7 | 249.5 |
|  | SD | 0.16 | 0.01 | 0.01 | 12.9 | 0.00 | 9.1 | 3.8 | 7.5 |
| MP+N+ | mean | 5.18 | 0.83 | 0.82 | 117.9 | 0.10 | 12.9 | 163.5 | 270.7 |
|  | SD | 0.07 | 0.01 | 0.01 | 48.1 | 0.00 | 6.7 | 13.9 | 20.3 |

DM dry matter, TC total carbon, TOC total organic carbon, C<sub>CWC</sub> cold water soluble carbon, N<sub>t</sub> total, N<sub>CWN</sub> cold water soluble nitrogen

**Table 4.**
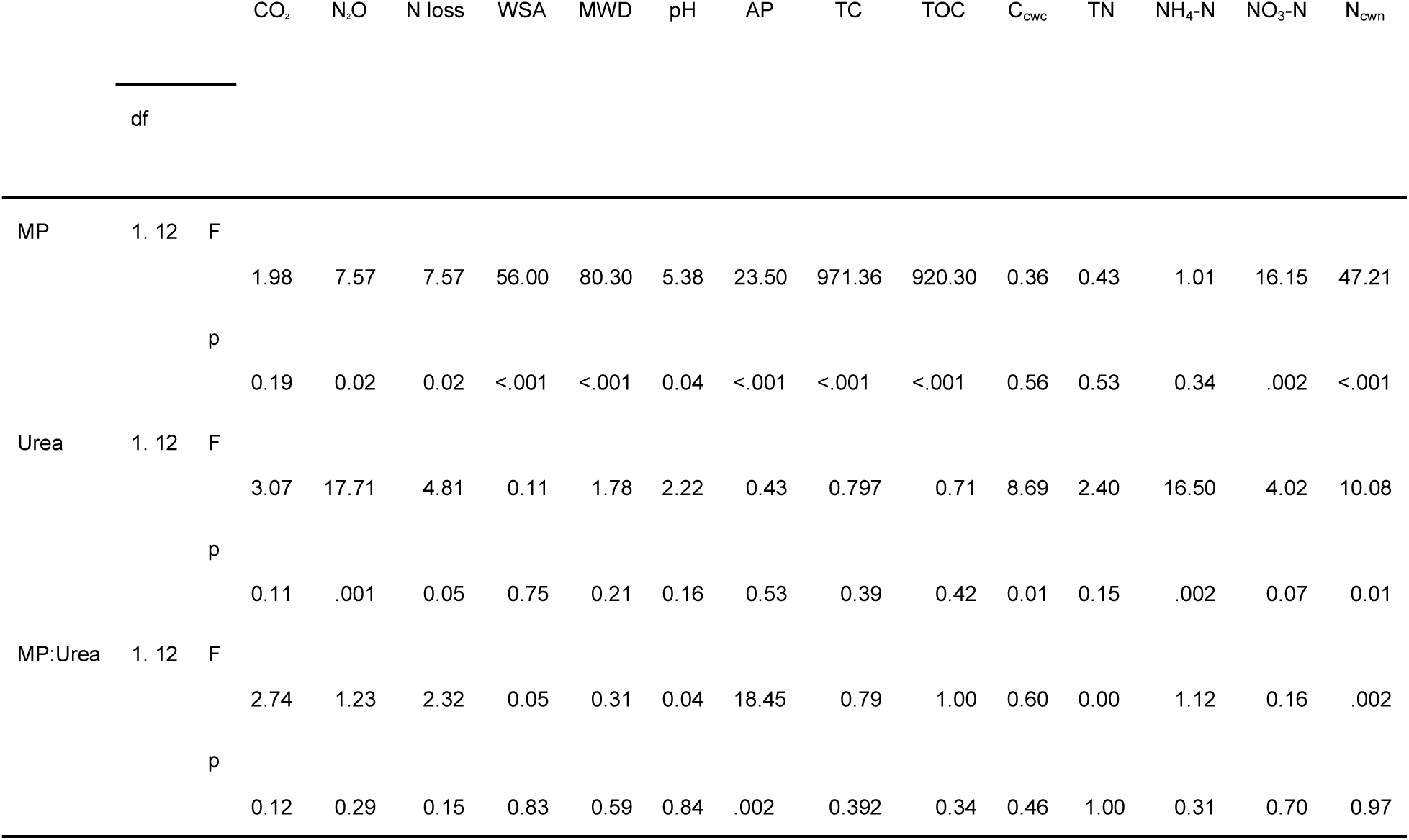
Model outcomes (n=16; with n=13 for AP) for the last harvest day (day 40) for CO_2_ emissions (in mg CO_2_-C core^−1^), N_2_O emissions (in mg N_2_O-N core^−1^), fertilizer N loss (in % of applied N), water-stability of soil aggregates (WSA, in %), mean weight diameter (MWD, in mm), and soil chemical characteristics (pH, AP, TC, TOC, C_cwc_, TN, NH_4_-N, NO_3_-N and N_cwn_).

The analysis of the relationships between soil properties and the cumulative CO_2_ and N_2_O fluxes showed that in the case of CO_2_ only closer relationships could be detected to the ammonium N content of the soil and in the case of N_2_O to the ammonium N and cold-water soluble carbon content of the soil (Figure 3).

**Figure 3.**
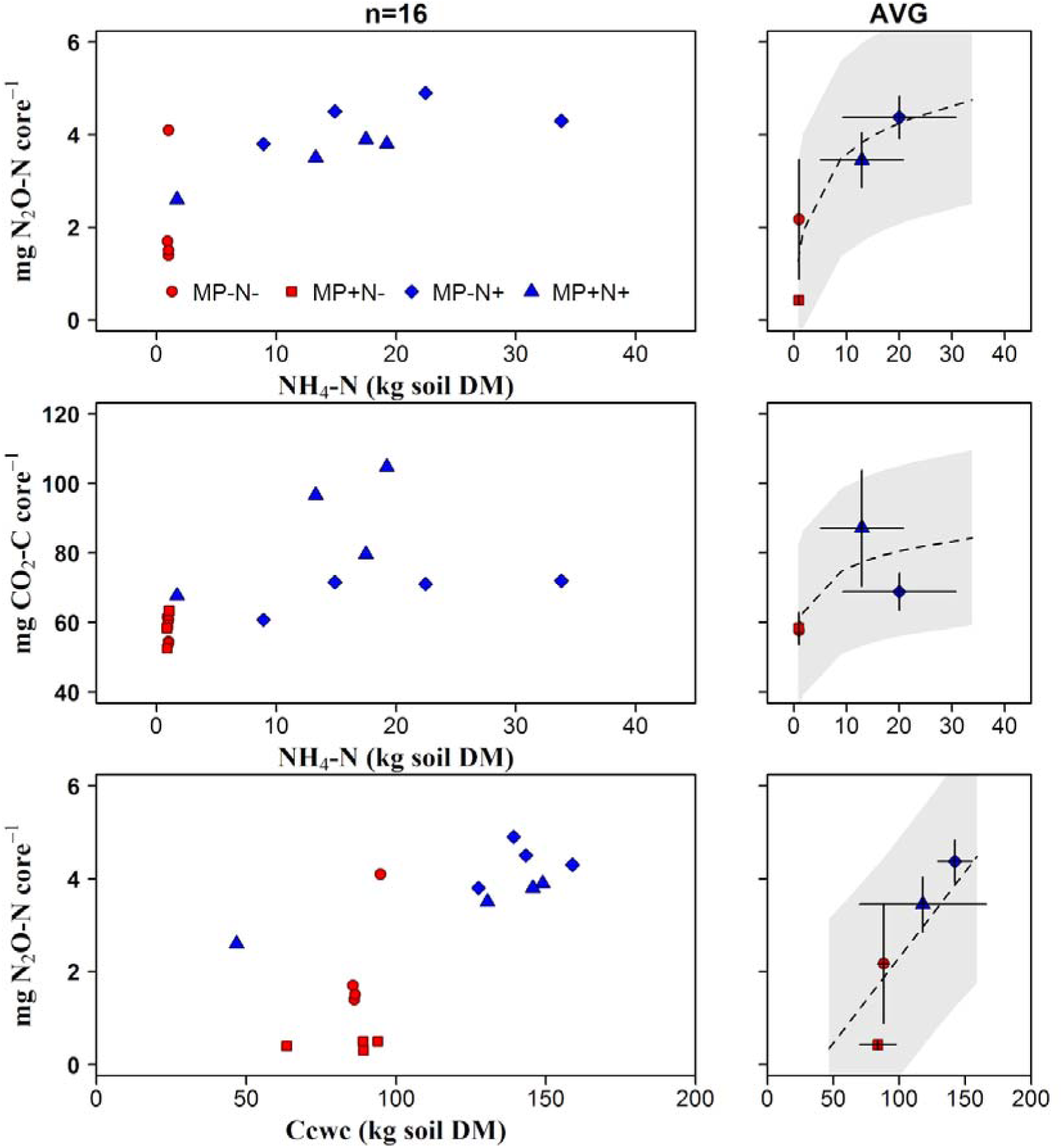
(a) Relationship between ammonium content of the soil and cumulative N_2_O emission; (b) Relationship between ammonium content of the soil and cumulative CO_2_ emission; (c) Relationship between content of cold water extractable soil C and cumulative N_2_O emission. (AVG = average per treatments combination). Treatment combinations are indicated by different colors (red = no N; blue with N added) and symbols.

## Discussion

We here present clear evidence that microplastic fibers affect the dynamics and intensity of trace gas fluxes, in particular of CO_2_ and N_2_O, from a sandy, intensively fertilized agricultural loam soil. It is interesting that microplastics mitigate the promotion of N_2_O release during intensive N fertilization. In the following we discuss these findings and the degree to which they are generalizable.

### Experimental approach allows clear determination of the effect of the test factors

In the first phase of the experiment, a clear effect of microplastic on gas flux dynamics was also observed, but over a period of about 18 days these effects were largely masked by the type of experimental approach, i.e. rewetting of the dry soil. This was probably due to a temporarily increased supply of microbially easily degradable C and N compounds as a result of soil disturbance and the addition of the diluted biogas digestate. By applying the urea only after this phenomenon had subsided, it was possible in the second phase of the experiment to clearly separate the effects of the test factors microplastics and N-fertilization. The renewed increase in CO_2_ and N_2_O flux rates is certainly due to the N fertilization and partially to the slight increase in WFPS to the final value of 55% (Fig. 3).

### Impact of N fertilization on CO_2_ and N_2_O fluxes

The applied N-fertilizers biogas digestate and urea both contained microbially easily degradable nitrogen and carbon. As expected, this led to a significant increase in both CO_2_ and N_2_O emissions. The effect of fertilization exceeded that of soil contamination with microplastics in both phases of the experiment. The investigations of Ren et al. ^28^ led to very similar results. Sources for the CO_2_ and N_2_O formation could have been the fertilizers themselves as well as an increased mineralization of soil organic matter induced by them as a result of the priming effect ^58, 59^.

However, it should be emphasized that the presence of microplastics in the simulated intensive N-fertilization arable soil led to a significant reduction not only of N_2_O emissions but also of fertilizer-derived N losses (Table 3). In view of this, it would be useful to examine in subsequent studies whether practical measures for reducing N_2_O emissions in agriculture can be derived from findings on the underlying mechanisms connecting with the impact of microplastics on soil structure.

### Evidence that the effect of microplastics on greenhouse gas fluxes is mainly due to changes in soil structure

The following facts indicate that the addition of microplastics has influenced the emissions of CO_2_ and N_2_O mainly through changes in soil structure. (i) The simultaneous change in the proportion of water-soluble aggregates and the gas flows in the variants with microplastics. (ii) Preventing the increase in N_2_O emission after increasing the water-filled pore space in the second phase of incubation for the variants not fertilized with urea (Figure 2, treatment MP-N-vs. MP+N-).

However, these findings do not provide information exactly which parameters of soil structure caused the changed CO_2_ and N_2_O fluxes. This requires further investigations in which parameters such as relative gas diffusivity, air permeability, air connectivity, air distance, air tortuosity, which allow a clear quantification of the gas movement in the soil, are systematically determined ^39, 60, 61^.

### Oxygen supply of the soil is probably the reason for the different reaction of CO_2_ and N_2_O on microplastics

There is substantial evidence that the addition of microplastics influenced the gas fluxes, mediated by the soil structure, in fact mainly by a changed oxygen supply. This also offers a clear explanation of the diverging effects for CO_2_ and N_2_O. Microplastic addition likely improved soil aeration: we showed an increase in soil aggregation and air permeability. This could explain the increased formation of CO_2_ because of mineralization proceeding more effectively in the presence of increased oxygen supply ^34, 44, 45, 62^.

The reduction of N_2_O emission after the addition of microplastics should also be due to a changed O_2_ supply. However, more complex processes are involved here than in CO_2_ release. A wide range of pathways can be involved in the formation and consumption of N_2_O. The most important of these are autotrophic and heterotrophic nitrification, nitrifier denitrification, heterotrophic denitrification, anaerobic ammonium oxidation (anammox) and dissimilatory nitrate reduction to ammonium (DNRA, or nitrate ammonification). In connection with our investigations, it is particularly important that N_2_O formation and N_2_O consumption can occur both in the presence and absence of O_2_ ^11, 63, 64^. In our study, nitrous oxide formation appears to be due to both autotrophic nitrification and heterotrophic denitrification. Some lines of evidence point to this. Previous investigations show that the WFPS we set up in the second phase of the investigation is almost optimal for the nitrification process taking place under aerobic conditions ^33, 62, 65-67^. This is also indicated by the above-mentioned increase in N_2_O release after raising the WFPS from 48 to 55% in the second phase of incubation (Figure 2, treatment MP-N-). Moreover, the close correlation between ammonium content of the soil and cumulative N_2_O emission indicates that autotrophic nitrification plays a significant role (Figure 3). However, the occurrence of microanaerobic zones, especially in highly aggregated soils, means that even at average WFPF values, part of the N_2_O can be produced in the course of denitrification and other processes that only occur in the absence of O_2_ ^37, 42, 43, 65, 66^. Therefore, the observed reduction in N_2_O release in the presence of microplastic, presumably due to increased O_2_ replenishment, can be regarded as an indication of reduced denitrification (Table 2). To find out if this is the case and to elucidate the real proportion of various processes contributing to N2O release, further investigations are required, especially on the basis of isotopic approaches in combination with quantification of gas movement into the soil ^33, 65, 66^.

### Recommendations for a more comprehensive assessment of the effect of microplastics on greenhouse gas fluxes

Our monitoring approach entailed high-frequency measurements; this is necessary to obtain precise results for trace gas emissions ^28, 68-72^. Future endeavors aimed at quantifying trace gas flux responses to microplastic addition should also rely on such measurements.

We carried out this experiment to study effects of microplastic fibers on soil under highly controlled conditions, excluding the role of plant roots or larger soil animals such as earthworms. In agricultural systems, plants may modify dynamics and trace gas fluxes. It is not clear which direction such modifications would take, because plants can affect outcomes in complicated ways in terms of their effects on rhizodeposition, competition for N, or changes in soil moisture ^73-77^. It is thus a high priority to include plant responses in assessments of trace gas fluxes when soils are exposed to microplastic.

Our study used microplastic fibers, which is a common shape of microplastics in the environment, but microplastics come in a wide variety of shapes ^50^, chemistries, and with many different additives present in consumer products ^78^. The only other experiment to test greenhouse gas effects used PE particles with a much higher concentration than in our investigations ^28^. Other shapes to examine include films, which have been shown to affect formation of cracks and water fluxes ^79^; this is relevant in agricultural systems owing to the prevalent use of mulching films. Different chemistries, including non-intentional additives and other compounds, may also affect different microbial players in the nutrient cycles leading to greenhouse gas emissions ^28^, possibly also leading to effects diverging from the ones observed here. Clearly, examining a broader parameter space of microplastic properties should be a priority for future research.

## Conclusions

Our laboratory study has clearly shown that microplastic fibers can influence trace gas emissions, and that soil structure effects are key to understanding such responses. Many studies of microplastic focus on a more classical ecotoxicological perspective, but our results suggest that microplastic should not be ignored in future estimates of greenhouse gas emissions and in assessing the actual risk to the environment from excessive N fertilization. Given the widespread presence of microplastic, especially in agricultural fields, such findings are relevant for understanding potential Earth system feedbacks of microplastic contamination ^14^. It is clear that ecosystem-level feedbacks should be included as well to achieve a more complete assessment of impacts.

## Data availability statement

Data supporting the findings of this study are freely accessible at the ZALF open-research-data repository (http://doi.org/104228/ZALF.DK.152).

## Acknowledgements

MCR acknowledges funding from a European Research Council Advanced Grant (694368), and the BMBF-funded project µPlastic (031B0907A). JA acknowledges funding through the Fachagentur Nachwachsende Rohstoffe (FNR) project ‘Krumensenke’. We thank Marten Schmidt and Bertram Gusovius for construction and for maintenance of the measurement apparatus.

## Supplementary Information

**Figure S1.**
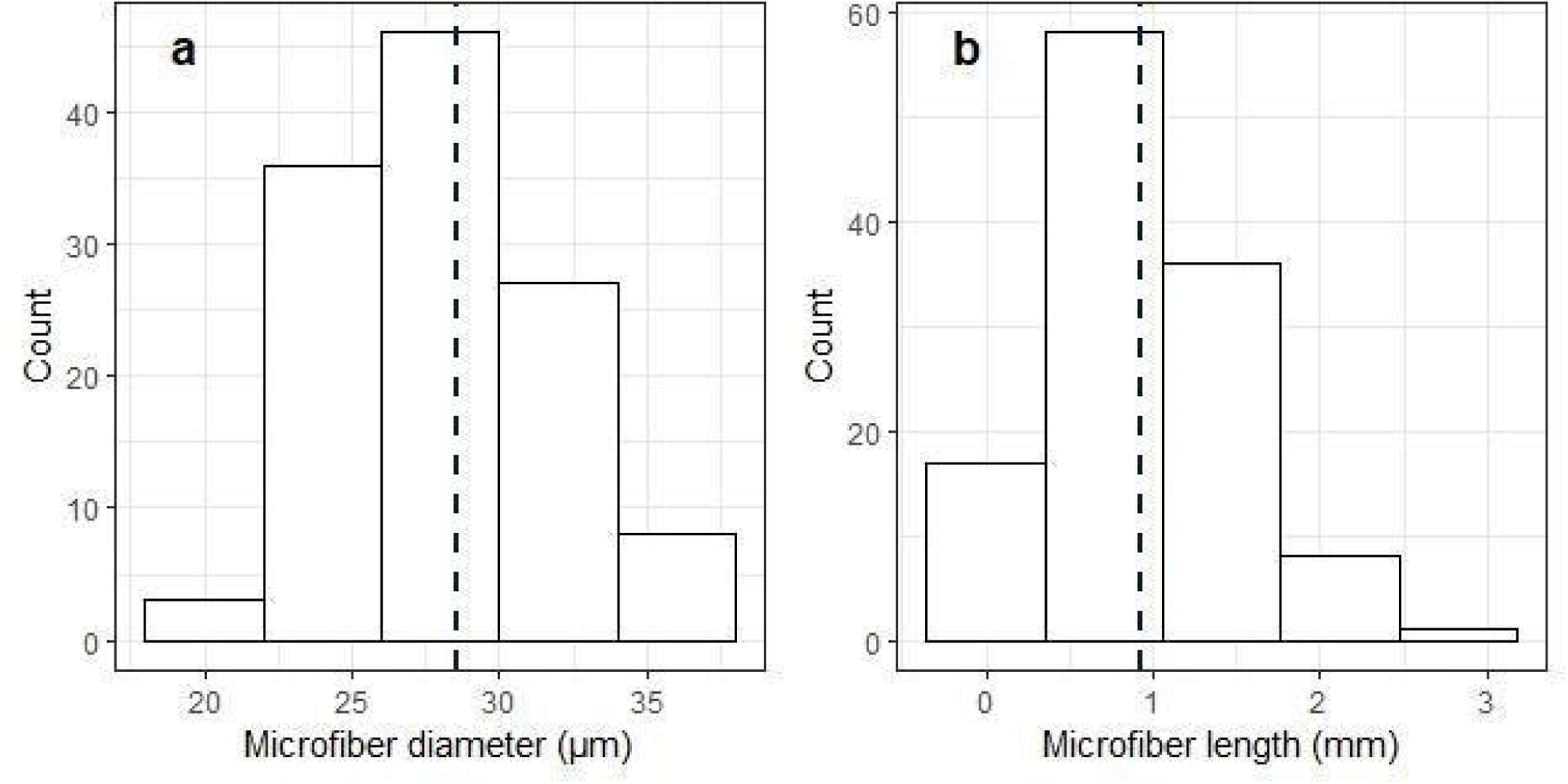
Diameter and length distribution of the microplastic fibers (n=120) used in this study. Data are based on measurements on pictures of fibers taken with a binocular microscope using image analysis with ImageJ.

